# Phylogenomic Characterization of Two *Alternaria* Lineages Associated with Cotton Leaf Spot in Tennessee

**DOI:** 10.64898/2026.10.17.759828

**Authors:** Ari Mortensen, Sahar Salimi, Brandon Hein, Riley Putman, Jinfa Zhang, Mostafa Rahnama

## Abstract

Alternaria leaf spot is an important foliar disease of cotton (*Gossypium hirsutum* L.) and has historically been associated primarily with *Alternaria macrospora* in the United States. In November 2025, symptomatic Upland cotton leaves from a research greenhouse at Tennessee Technological University’s Oakley Farm in Livingston, Tennessee, yielded two *Alternaria* isolates, 2-2 and 9-1. Quantitative conidial characterization showed substantial overlap in dimensions, septation, and shape. Concatenated ITS-*RPB2* phylogenetic analysis supported different broad affinities, placing isolate 2-2 within an *A. alternata*-dominated group and isolate 9-1 within a group associated with the *A. arborescens* species complex (AASC), although their immediate relationships were weakly resolved. Phylogenomic analysis of 1,501 single-copy BUSCO orthologs from 63 genomes placed isolate 2-2 within the *A. alternata* lineage, sister to BMP2343, and isolate 9-1 within the AASC, sister to MOD1-FUNGI6. In controlled pathogenicity assays, isolate 9-1 reproduced foliar lesions on cotton, whereas mock-inoculated plants remained symptomless, and molecularly confirmed re-isolation fulfilled Koch’s postulates. Isolate 2-2 produced insufficient conidia under the culture conditions used to prepare a standardized inoculum, and its pathogenicity was therefore not determined. These results reveal phylogenetically distinct *Alternaria* lineages associated with symptomatic cotton and establish an AASC isolate as a causal agent of cotton leaf spot in Tennessee.

---

Cotton (*Gossypium* spp.) is one of the world’s most important fiber crops, with Upland cotton (*Gossypium hirsutum* L.) accounting for approximately 95% of global cotton fiber production (Liu et al. 2020; Stewart 2025). Alternaria leaf spot, also referred to as Alternaria leaf blight, occurs in cotton-growing regions worldwide and has historically been associated primarily with *Alternaria macrospora* Zimm. and *A. alternata* (Fr.) Keissl. (Bashan et al. 1991; Zhu et al. 2019). Although A. macrospora has long been regarded as the principal cotton-associated species, accumulating evidence indicates that A. alternata can also act as a primary cotton pathogen (Le and Gregson 2019; Mohamed et al. 2019; Zhu et al. 2019), with the relative importance of these taxa varying among geographic regions.

Several studies have demonstrated the pathogenic role of *A. alternata* on cotton. In southern New Mexico, surveys of symptomatic commercial cotton fields recovered *A. alternata* from Alternaria leaf spot lesions, and pathogenicity assays confirmed its ability to reproduce the disease on cotton; notably, *A. macrospora*, historically considered the principal causal agent, was not recovered from the representative isolates examined (Zhu et al. 2019). *A. alternata* was likewise the predominant *Alternaria* species associated with Alternaria leaf spot on cotton seedlings in southern New South Wales, Australia, where molecular identification showed that most recovered isolates belonged to *A. alternata* (Le and Gregson 2019). *A. alternata* has also been confirmed as a causal agent of cotton leaf blight in Sudan (Mohamed et al. 2019), and its association with Alternaria leaf spot has been documented in other cotton-growing regions including Turkey (Olmez et al. 2023). Collectively, these studies indicate that the etiology of cotton *Alternaria* leaf spot is more diverse than historically recognized.

Despite the documented importance of *A. alternata* on cotton in other production regions, the identity of *Alternaria* associated with cotton leaf spot in Tennessee remains poorly characterized. The available published Tennessee record comes from field experiments conducted at Jackson , western Tennessee, during the 2008-2010 growing seasons, in which foliar disease on Upland cotton was attributed to *A. macrospora* (Woodward et al. 2016). Thus, it remains unknown whether other *Alternaria* lineages, including those associated with cotton leaf spot elsewhere, can also cause disease on cotton in Tennessee. Determining the identity and pathogenicity of such lineages is important for defining the etiological diversity of Alternaria leaf spot in the state.

Accurate identification of cotton-associated *Alternaria* is complicated by the close evolutionary relationships among small-spored taxa within *Alternaria* sect. *Alternaria*. Morphological characteristics can overlap substantially, and commonly used barcode loci may provide limited phylogenetic resolution, contributing to extensive taxonomic reassessment within the section (Woudenberg et al. 2013, 2015). Previous studies of cotton-associated *Alternaria* have therefore combined morphology and multiple DNA markers for identification (Le and Gregson 2019; Zhu et al. 2019). Whole-genome data provide an additional level of resolution for placing isolates relative to characterized lineages. Here, we combined quantitative conidial morphology, ITS-*RPB2* phylogenetics, pathogenicity testing, and whole-genome phylogenomics to characterize two *Alternaria* isolates recovered from symptomatic cotton in Tennessee, resolve their phylogenetic placement, and evaluate pathogenicity using a standardized conidial inoculation assay.

In November 2025, leaves of Upland cotton (*Gossypium hirsutum* L.) exhibiting dark-brown to black necrotic spots of variable size, some with tan centers, were collected from plants grown in a research greenhouse at Tennessee Technological University’s Oakley Farm in Livingston, Tennessee, USA. Lesions occasionally coalesced into larger irregular areas and were associated with localized chlorosis (Fig. 1A). Isolates 2-2 and 9-1 originated from separate symptomatic leaves displaying similar leaf-spot symptoms. Approximately 3 cm² of symptomatic leaf tissue was excised, rinsed to remove surface debris, immersed in 3% NaOCl for 1 min, and rinsed sequentially in sterile water. Surface-disinfested tissues were cut into approximately 2-mm² fragments, placed directly on potato dextrose agar (PDA), and incubated at 25 to 27°C for 2 to 3 days. Emerging colonies were transferred repeatedly to fresh medium to obtain purified cultures. Because bacterial contamination was observed in some initial cultures, subsequent purification included PDA amended with 100 µg/ml ampicillin. Eight initial fungal cultures were recovered, and repeated subculturing and microscopic examination identified cultures with *Alternaria*-like conidia. As a result, two representative isolates, designated 2-2 and 9-1, were selected for further morphological, molecular, and genomic characterization.

**Fig. 1.**
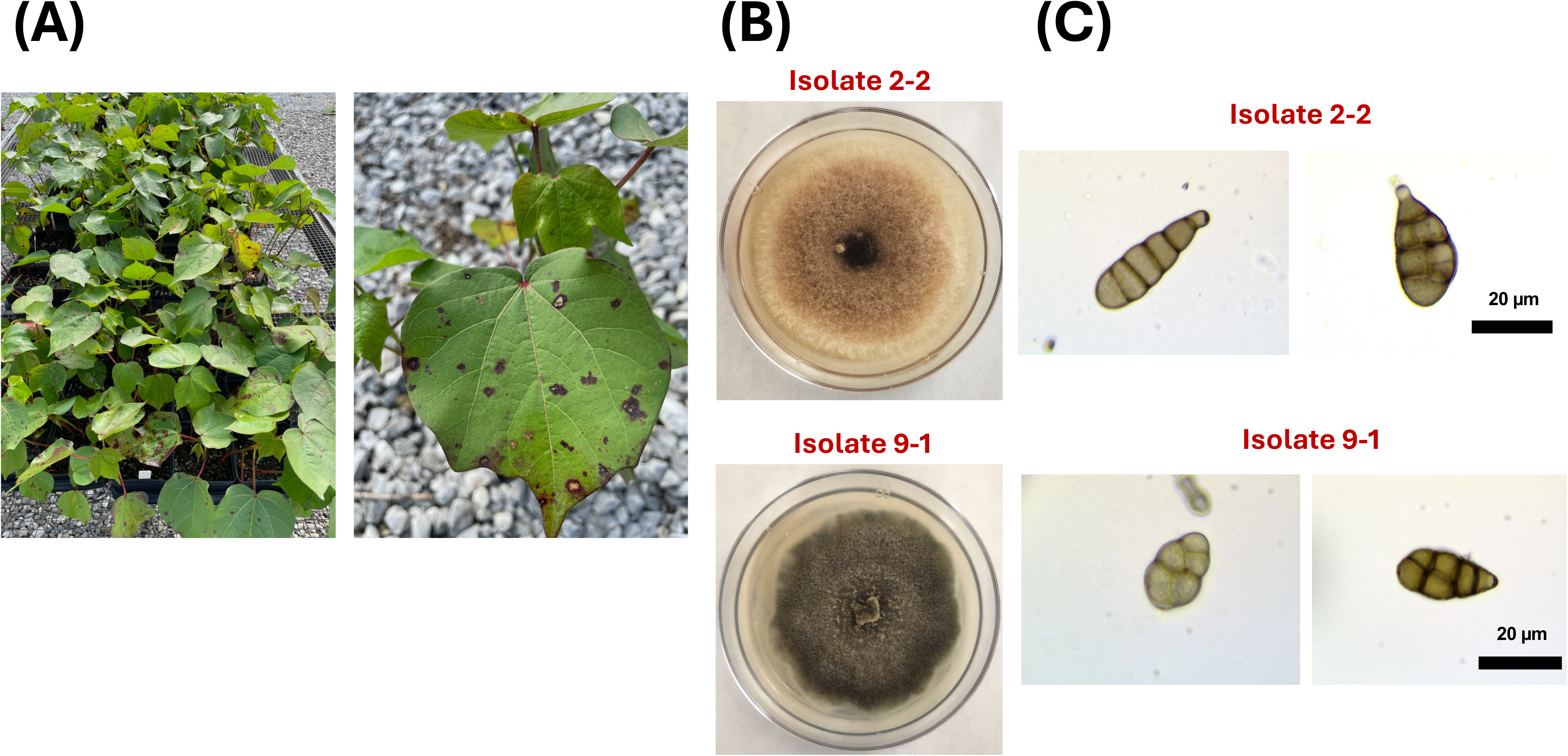
Natural symptoms, colony morphology, and conidial morphology of cotton-associated *Alternaria* isolates 2-2 and 9-1 from Tennessee. (A) Cotton plants exhibiting naturally occurring leaf-spot symptoms and a representative symptomatic leaf from which *Alternaria* isolates were recovered. (B) Representative colony morphology of isolates 2-2 (top) and 9-1 (bottom) on potato dextrose agar. (C) Representative conidia of isolate 2-2 (top) and isolate 9-1 (bottom), illustrating the range of conidial morphologies observed; catenulate conidia are shown for isolate 9-1. Scale bars = 20 µm.

Initial microscopic observations revealed septate conidia consistent with *Alternaria*. Conidial morphology was characterized quantitatively following a general fungal conidial measurement approach (Szarka et al. 2023). Conidia were examined from PDA cultures grown at 25°C between 3 and 11 days after plate inoculation (DPI). In total, 31 mature, intact, and clearly resolved conidia of isolate 2-2 and 30 conidia of isolate 9-1 were examined. Conidia were imaged using an Olympus AX70/Provis microscope equipped with an AmScope MU1000B digital camera at either 400× or 1,000× total magnification. Conidial dimensions were measured using ImageJ (Schneider et al. 2012) after independent calibration of the 40× and 100× objectives using the 10-µm graduations of a 1-mm calibration slide supplied with the AmScope microscope. For each conidium, total length, maximum width, number of transverse septa, number of longitudinal or oblique septa, presence of partial septa, overall shape, and arrangement as solitary or catenulate were recorded. Shape was classified as elongate/obclavate, ellipsoid, ovoid/obovoid, or irregular/broad. Tiny developing terminal cells, damaged or collapsed conidia, overlapping structures, and conidia for which boundaries or septa could not be reliably resolved were excluded. Length and width are reported as range and mean ± standard deviation, whereas septation is reported as the observed range and conidial shape and arrangement as frequencies or percentages. Individual measurements and imaging magnification are provided in Supplementary Table S1.

Conidia of isolate 2-2 measured 10.54 to 42.04 × 6.94 to 18.51 µm (mean ± SD, 26.12 ± 9.12 × 12.13 ± 3.13 µm; n = 31), with 1 to 5 transverse and 0 to 5 longitudinal septa. Conidia of isolate 9-1 measured 12.21 to 37.42 × 7.24 to 18.80 µm (mean ± SD, 23.09 ± 6.04 × 11.46 ± 2.68 µm; n = 30), with 1 to 5 transverse and 0 to 4 longitudinal septa. Conidial shapes overlapped substantially between the two isolates. In isolate 2-2, 14 conidia (45.2%) were ovoid/obovoid, 13 (41.9%) elongate/obclavate, 2 (6.5%) ellipsoid, and 2 (6.5%) irregular/broad. In isolate 9 -1, 13 conidia (43.3%) were elongate/obclavate, 11 (36.7%) ovoid/obovoid, 4 (13.3%) ellipsoid, and 2 (6.7%) irregular/broad. All conidia of isolate 2-2 were observed singly, whereas 27 (90.0%) conidia of isolate 9-1 were solitary and 3 (10.0%) were observed in chains. Overall, conidial dimensions, septation, and shape showed substantial overlap between the two isolates (Supplementary Table S1). Representative colony and conidial morphologies are shown in Fig. 1B and C, respectively. Because conidial morphology did not clearly distinguish the two isolates, their phylogenetic affinities were next evaluated using ITS and *RPB2*.

Genomic DNA was extracted from fungal mycelium after cultures were grown for 5 days in potato dextrose broth at 25°C with shaking at 150 rpm, as described by Rahnama et al. (2020). Initial molecular characterization was based on the nuclear ribosomal internal transcribed spacer region (ITS) and the second-largest subunit of RNA polymerase II (*RPB2*). The ITS region was amplified using primers ITS5 (5′-GGAAGTAAAAGTCGTAACAAGG-3′) and ITS4 (5′-TCCTCCGCTTATTGATATGC-3′) (White et al. 1990), whereas *RPB2* was amplified using primers fRPB2-5F2 (5′-GGGGWGAYCAGAAGAAGGC-3′; Sung et al. 2007) and fRPB2-7cR (5′-CCCATRGCTTGYTTRCCCAT-3′; Liu et al. 1999). PCR amplification was performed using Q5 High-Fidelity 2× Master Mix (New England Biolabs, Ipswich, MA, USA). Following the manufacturer’s standard reaction setup, 25-µl reactions contained 12.5 µl of Q5 2× Master Mix, 1.25 µl each of 10 µM forward and reverse primers (0.5 µM final concentration), template DNA, and nuclease-free water to 25 µl. PCR conditions were as follows: for *RPB2*, 32 cycles were performed with annealing at 64°C for 30 s; for ITS, a 32-cycle touchdown program was used, with the annealing temperature decreasing from 72°C to 62°C during the first 20 cycles, followed by 12 cycles at 62°C. All other cycling conditions followed the manufacturer’s standard Q5 protocol, including initial denaturation at 98°C for 30 s, denaturation at 98°C for 10 s, extension at 72°C for 30 s, and a final extension at 72°C for 2 min. PCR products were visualized by agarose gel electrophoresis, and products requiring separation from nonspecific amplification were excised and gel purified. Purified ITS and *RPB2* products from isolates 2-2 and 9-1 were submitted to MCLAB (South San Francisco, CA, U.S.A.) for bidirectional Sanger sequencing using the corresponding amplification primers. To assess their phylogenetic placement, concatenated ITS and *RPB2* sequences from isolates 2-2 and 9-1 were analyzed together with 163 reference isolates selected primarily from Woudenberg et al. (2015), with *A. alternantherae* CBS 124392 included as the outgroup for tree visualization (Supplementary Table S2).

ITS and *RPB2* sequences were aligned using Clustal Omega v1.2.3 (Sievers et al. 2011), as implemented in Geneious Prime 2025.2.1 (Biomatters Ltd., Auckland, New Zealand), and concatenated into a final alignment comprising 165 taxa and 1,279 nucleotide positions, of which 68 were parsimony-informative. Maximum-likelihood phylogenetic inference was performed using IQ-TREE v2.2.2 (Minh et al. 2020). ModelFinder (Kalyaanamoorthy et al. 2017) selected TNe+G4 as the best-fitting nucleotide substitution model according to the Bayesian information criterion, and branch support was assessed using 1,000 ultrafast bootstrap replicates (Hoang et al. 2018). The maximum-likelihood tree was rooted for visualization with *A. alternantherae* CBS 124392 and is presented in Supplementary Fig. S1.

The concatenated ITS-*RPB2* analysis supported different broad phylogenetic affinities for isolates 2-2 and 9-1 but showed limited resolution of relationships within sect. *Alternaria* (Supplementary Fig. S1). Isolate 2-2 occurred within an *A. alternata*-dominated clade; although the branch immediately associating 2-2 with the surrounding reference cluster was weakly supported (UFBoot = 23), a broader *A. alternata*-dominated clade containing 2-2 received UFBoot support of 96. In contrast, isolate 9-1 occurred within a portion of the tree associated with the *Alternaria arborescens* E.G. Simmons species complex (AASC). Its immediate placement was also weakly supported (UFBoot = 14), whereas a broader clade containing 9-1 together with *A. arborescens* and *A. aff. arborescens* reference isolates received UFBoot support of 97. Thus, the two-locus analysis was consistent with different phylogenetic affinities of the two cotton isolates but lacked sufficient local resolution for definitive lineage assignment. Definitive placement was therefore based on the whole-genome phylogenomic analysis described below. The resulting ITS and *RPB2* sequences were deposited in GenBank under accession numbers PZ923877 and PZ944210 for isolate 2-2 and PZ923876 and PZ944211 for isolate 9-1, respectively.

To evaluate pathogenicity, a standardized conidial inoculation assay was attempted for both isolates using growth chamber-grown cotton plants and a protocol adapted from Zhu et al. (2019). Although isolate 2-2 produced conidia sufficient for morphological characterization, conidial production under the culture conditions used was insufficient to prepare the standardized inoculum required for pathogenicity testing. Pathogenicity testing was therefore completed for isolate 9-1 only. Upland cotton (*G. hirsutum*) plants were grown four plants per pot in commercial garden soil containing fertilizer under a 12-h light/12-h dark photoperiod at 30°C/25°C, within the range of growth conditions used in previous cotton–*Alternaria* studies (Bashan 1984; Zhu et al. 2019), with trays of water placed in the chamber to increase humidity. Conidia of isolate 9-1 were harvested from sporulating PDA cultures into sterile water, pooled, and quantified using a hemocytometer. Ten milliliters of conidial suspension (approximately 5 × 10^5 conidia/ml) was sprayed onto the foliage of inoculated plants, whereas mock-inoculated controls received an equal volume of sterile Milli-Q water. Three replicate pots, each containing four plants, were used for the isolate 9-1 treatment and the mock control. Immediately after inoculation, plants were enclosed in plastic bags containing water to maintain high humidity and incubated at 25°C in darkness for the first 24 h, consistent with high-humidity postinoculation procedures used in previous cotton–*Alternaria* assays (Zhao et al. 2012; Zhu et al. 2019). Plants remained enclosed for approximately 48 h before being returned to the standard growth-chamber conditions and were subsequently misted with sterile water two to three times daily. Plants were monitored daily for development of foliar lesions.

To complete Koch’s postulates for isolate 9-1, fungi were re-isolated from experimentally induced lesions and molecularly compared with the inoculated isolate. At 7 days after inoculation, plants inoculated with isolate 9-1 exhibited foliar lesions, whereas mock-inoculated plants remained symptomless. Representative lesions were excised, rinsed in deionized water, surface disinfested in 3% NaOCl for 1 min, rinsed three times in sterile Milli-Q water, cut into approximately 2-mm² fragments, and placed on PDA supplemented with 100 µg/ml ampicillin. Plates were incubated at 25 to 27°C in darkness for 2 to 3 days, and representative fungal colonies recovered from symptomatic tissue were transferred to potato dextrose broth and processed for genomic DNA extraction as described above. Re-isolates were evaluated using the same ITS5/ITS4 and fRPB2-5F2/fRPB2-7cR amplification and Sanger sequencing procedures used for the original isolate. Representative re-isolates recovered from plants inoculated with 9-1 matched the original isolate at both ITS and *RPB2*. *Alternaria* was not recovered from mock-inoculated controls. Together, these results fulfilled Koch’s postulates for isolate 9-1.

Draft genome assemblies of isolates 2-2 and 9-1 were generated from Illumina NextSeq 1000 2 × 100-bp paired-end reads. The resulting assemblies were 33.17 and 33.72 Mb, respectively, and were deposited in DDBJ/ENA/GenBank under accession numbers JCDBFJ000000000 (2-2) and JCDBFI000000000 (9-1). These assemblies were used for the phylogenomic analyses described below.

To resolve the genome-scale phylogenetic placement of isolates 2-2 and 9-1, their assemblies were analyzed together with 61 publicly available *Alternaria* reference genomes, producing a 63-taxon data set (Fig. 2; Supplementary Table S3). Reference genomes represented the major lineages sampled within sect. *Alternaria*, including the *A. alternata* lineage, AASC, *A. gaisen*, *A. longipes*, *A. burnsii*, and additional historically named or currently unresolved lineages. *Alternaria macrospora* BMP1949 and *A. solani* HWC-168-2012p from sect. *Porri* were included as outgroup taxa. Reference assemblies were obtained from NCBI and JGI and were selected primarily from the genome collections of Kim and Dettman (2025) and Dettman et al. (2026).

**Fig. 2.**
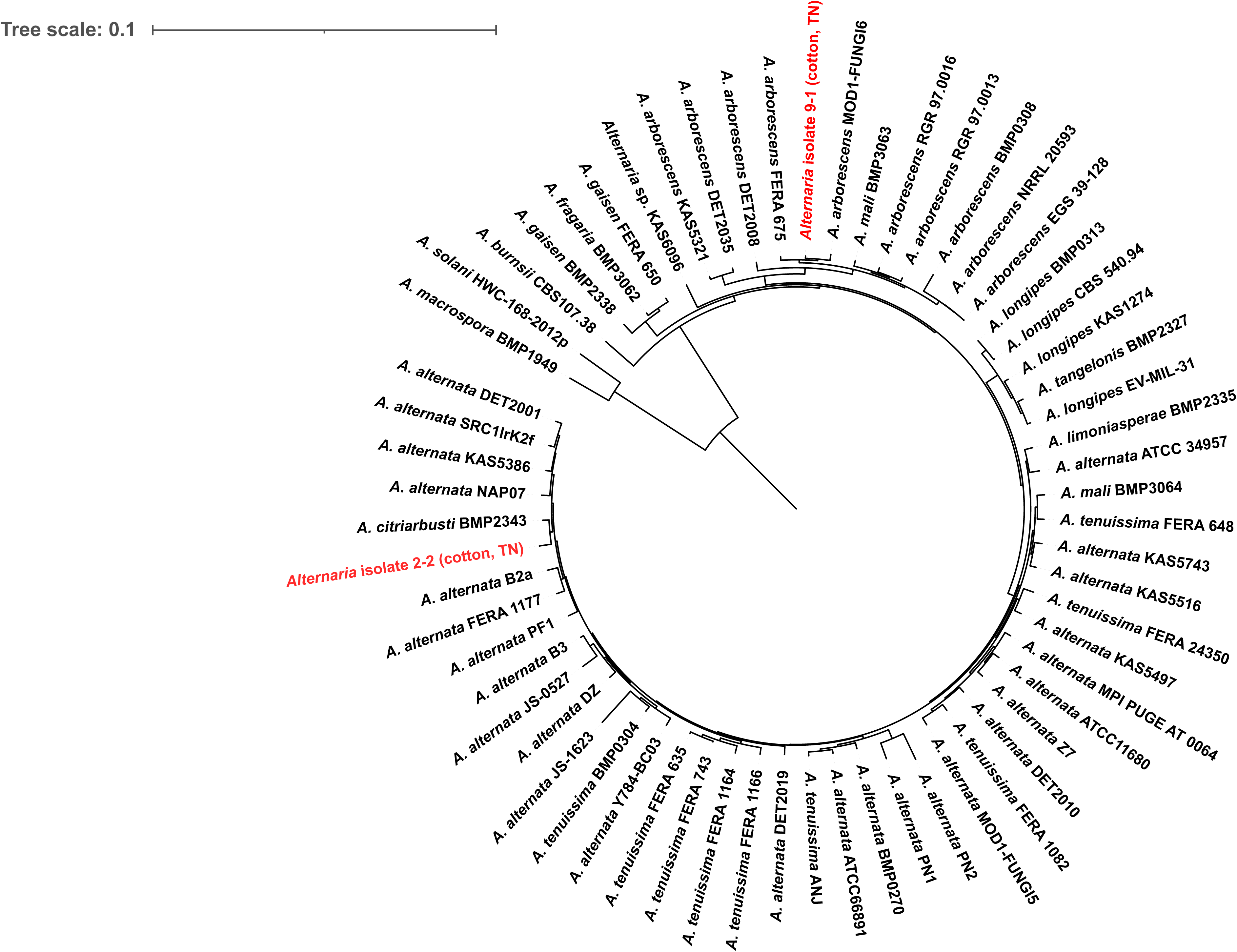
Maximum-likelihood phylogenomic tree of cotton-associated *Alternaria* isolates 2-2 and 9-1 and 61 publicly available reference genomes. The tree was inferred from a concatenated amino-acid alignment of 1,501 single-copy BUSCO orthologs present in ≥95% of the 63 taxa (1,024,950 aligned positions) using IQ-TREE v2.3.6 and displayed in circular format. ModelFinder with partition merging identified 52 merged partitions. The tree was rooted with the sect. *Porri* taxa *A. macrospora* BMP1949 and *A. solani* HWC-168-2012p. The Tennessee cotton isolates are highlighted in red. UFBoot support values are omitted for clarity and are shown on the corresponding rectangular tree in Supplementary Fig. S2.

Single-copy orthologs were identified with BUSCO v5.4.3 (Manni et al. 2021) using the ascomycota_odb10 lineage data set. BUSCO loci present as single-copy orthologs in at least 95% of the 63 taxa were retained, resulting in 1,501 loci. Amino-acid sequences for each locus were aligned independently using MAFFT v7.505 (Katoh and Standley 2013) and trimmed using ClipKIT v2.14.0 (Steenwyk et al. 2020) with the smart-gap option. The resulting alignments were concatenated into a supermatrix containing 1,024,950 amino-acid positions. Maximum-likelihood phylogenetic inference was performed with IQ-TREE v2.3.6 (Minh et al. 2020) using an edge-linked proportional partition model (Chernomor et al. 2016). Each BUSCO locus was initially treated as an independent partition, and ModelFinder with partition merging (MFP+MERGE; Kalyaanamoorthy et al. 2017) identified an optimized scheme of 52 merged partitions. Branch support was assessed with 1,000 SH-like approximate likelihood ratio test (SH-aLRT) replicates (Guindon et al. 2010) and 1,000 ultrafast bootstrap (UFBoot) replicates (Hoang et al. 2018). The maximum-likelihood tree was rooted using the sect. *Porri* branch containing *A. macrospora* BMP1949 and *A. solani* HWC-168-2012p. The rooted tree was displayed in circular format using Interactive Tree Of Life (iTOL; Letunic and Bork 2024) for Fig. 2, whereas the corresponding rectangular support tree was visualized using ggtree (Yu et al. 2017) in R v4.4.0 (R Core Team 2024).

Consistent with the broad affinities observed in the ITS-*RPB2* analysis but providing substantially greater phylogenetic resolution, whole-genome phylogenomics clearly demonstrated that the two cotton isolates represent distinct evolutionary lineages (Fig. 2). Isolate 2-2 was placed within the *A. alternata* lineage and was recovered as sister to strain BMP2343, historically identified as *A. citriarbusti*, with 100% SH-aLRT and 100% UFBoot support. In contrast, isolate 9-1 was nested within the AASC and was recovered as sister to AASC strain MOD1-FUNGI6 with 100% SH-aLRT and 99% UFBoot support; this pair grouped with AASC strain FERA675 with 100% SH-aLRT and 99% UFBoot support. Thus, despite being recovered from cotton leaves displaying similar disease symptoms, isolates 2-2 and 9-1 represent two phylogenetically distinct lineages within *Alternaria* sect. *Alternaria*.

Together, the morphological, marker-based, and phylogenomic data demonstrate that two phylogenetically distinct *Alternaria* lineages were recovered from cotton leaves displaying similar leaf-spot symptoms. Isolate 2-2 belongs to the *A. alternata* lineage, whereas isolate 9-1 belongs to the AASC. Their conidial dimensions, septation, and shapes overlapped substantially, and ITS-*RPB2* data resolved only their broader affinities. In contrast, genome-scale phylogenetics clearly separated the two lineages, emphasizing the limitations of symptom phenotype, conidial morphology, and commonly used marker loci for lineage-level identification within *Alternaria* sect. *Alternaria*. Pathogenicity testing established isolate 9-1 as a causal agent of cotton leaf spot, with fulfillment of Koch’s postulates. The pathogenic role of isolate 2-2 remains unresolved because it did not produce sufficient conidia under the culture conditions used to prepare a standardized inoculum. Importantly, insufficient in vitro sporulation should not be interpreted as evidence that isolate 2-2 is nonpathogenic; rather, additional sporulation or inoculation approaches will be required to determine its relationship with cotton.

Alternaria leaf spot has previously been reported on cotton in Tennessee, where disease in field experiments at Jackson was attributed to *A. macrospora* (Woodward et al. 2016). Therefore, this study is not the first report of Alternaria leaf spot of cotton in Tennessee. To our knowledge, however, it is the first confirmed report of a member of the AASC causing cotton leaf spot in the state. The recovery of an *A. alternata*-lineage isolate from a separate symptomatic cotton leaf further demonstrates that phylogenetically distinct *Alternaria* lineages can occur on symptomatic cotton, although the pathogenic role of isolate 2-2 remains undetermined. Because both isolates originated from a research greenhouse rather than a systematic survey of commercial fields, their distribution and prevalence in Tennessee cotton production remain unknown. Surveys of commercial fields will be needed to determine their occurrence and agricultural significance.

## Acknowledgments

The authors thank Justin King for assistance with library preparation and Carla Hurt for assistance with Illumina sequencing using the Tennessee Tech University Department of Biology sequencing facility.

## GenBank accession numbers

ITS, PZ923877 (isolate 2-2) and PZ923876 (isolate 9-1); *RPB2*, PZ944210 (isolate 2-2) and PZ944211 (isolate 9-1); WGS, JCDBFJ000000000 (isolate 2-2) and JCDBFI000000000 (isolate 9-1); SRA, SRR40282443 (isolate 2-2) and SRR40282442 (isolate 9-1); BioProject PRJNA1514184.

## Data availability

ITS sequences generated in this study are available in GenBank under accession numbers PZ923877 for isolate 2-2 and PZ923876 for isolate 9-1. *RPB2* sequences are available under accession numbers PZ944210 for isolate 2-2 and PZ944211 for isolate 9-1. Raw Illumina reads are available under BioProject PRJNA1514184, with BioSamples SAMN62592141 (2-2) and SAMN62592142 (9-1) and SRA accessions SRR40282443 (2-2) and SRR40282442 (9-1). Draft genome assemblies are available in DDBJ/ENA/GenBank under accession numbers JCDBFJ000000000 for isolate 2-2 and JCDBFI000000000 for isolate 9-1.

## TABLES

**Supplementary Table S1. Quantitative conidial morphology of cotton-associated *Alternaria* isolates 2-2 and 9-1.** Conidia were examined from PDA cultures between 3 and 11 days after plate inoculation. A total of 31 mature, intact conidia of isolate 2-2 and 30 conidia of isolate 9-1 were measured from images acquired at 400× or 1,000× total magnification. Measurements included conidial length, maximum width, transverse septa, longitudinal septa, partial septa, shape, and arrangement. The Summary worksheet reports dimensional ranges and means ± standard deviations, septation ranges, and frequencies of conidial shape and arrangement; the Individual measurements worksheet provides the underlying measurements for each conidium.

**Supplementary Table S2. *Alternaria* isolates and sequence accessions included in the concatenated ITS-*RPB2* phylogenetic analysis.** The data set comprises cotton-associated isolates 2-2 and 9-1 and 163 reference isolates. For each isolate, the table provides the taxon name, strain or isolate designation, locality and host or substrate information, ITS and *RPB2* GenBank accession numbers, and source or role in the analysis. *Alternaria alternantherae* CBS 124392 was included as the outgroup for tree visualization.

**Supplementary Table S3. *Alternaria* genome assemblies included in the 63-taxon phylogenomic analysis.** The data set comprises cotton-associated isolates 2-2 and 9-1 and 61 publicly available reference genomes. For each genome, the table provides the taxon name, strain or isolate designation, lineage or section assignment used in this study, genome source, assembly accession or JGI genome identifier, and reference or source. *Alternaria macrospora* BMP1949 and *A. solani* HWC-168-2012p were included as sect. *Porri* outgroup taxa.

**Supplementary Fig. S1. Maximum-likelihood phylogenetic tree of cotton-associated *Alternaria* isolates 2-2 and 9-1 based on concatenated ITS and *RPB2* sequences.** The circular tree included 165 taxa and 1,279 aligned nucleotide positions, of which 68 were parsimony-informative. Maximum-likelihood inference was performed with IQ-TREE v2.2.2 under the TNe+G4 nucleotide substitution model selected by ModelFinder according to the Bayesian information criterion. Branch support was assessed using 1,000 ultrafast bootstrap (UFBoot) replicates, and all available UFBoot values are shown in red at internal nodes. The tree was rooted for visualization with *A. alternantherae* CBS 124392. *Alternaria* isolate 2-2 (cotton, TN) and *Alternaria* isolate 9-1 (cotton, TN) are highlighted in red.

**Supplementary Fig. S2. Maximum-likelihood phylogenomic tree of cotton-associated *Alternaria* isolates 2-2 and 9-1 and 61 publicly available reference genomes showing UFBoot support values.** The rooted rectangular tree represents the same topology and phylogenomic analysis shown in Fig. 2 and was based on 1,501 single-copy BUSCO orthologs present in ≥95% of the 63 taxa (1,024,950 aligned amino-acid positions). Branch support was assessed using 1,000 SH-like approximate likelihood ratio test (SH-aLRT) replicates and 1,000 ultrafast bootstrap (UFBoot) replicates; all available UFBoot values are shown at internal branches. The tree was rooted with the sect. *Porri* taxa *A. macrospora* BMP1949 and *A. solani* HWC-168-2012p. The Tennessee cotton isolates are highlighted in red.

